# HiC2Self: self-supervised denoising for bulk and single-cell Hi-C contact maps

**DOI:** 10.1101/2024.11.21.624767

**Authors:** Rui Yang, Alireza Karbalayghareh, Christina S. Leslie

## Abstract

Hi-C is a chromosome conformation capture assay used to study 3D genome organization. The recent development of single-cell Hi-C technologies has further enabled the examination of 3D chromatin organization in individual cells, although these approaches often suffer from low-coverage libraries and data sparsity. Here we introduce HiC2Self, a self-supervised framework for denoising Hi-C contact maps that requires only low-coverage data as input. HiC2Self not only reconstructs key structures (such as TADs and significant loops) from bulk libraries, but its self-supervised training framework also allows it to easily reconstruct cell-type-specific Hi-C structures without the generalization challenges faced by supervised models. HiC2Self can also accurately reconstruct significant loops from Micro-C data at 1 kb resolution. Moreover, when applied to single-nucleus methyl-3C data, HiC2Self successfully reconstructs local TAD structures around specific genes at 10 kb resolution with as few as 50 cells. Finally, HiC2Self enables the examination of single-cell structures at 50 kb resolution in individual cells of the same cell type. HiC2Self thus provides a general tool for denoising bulk, pseudo-bulk, and single-cell 3D contact maps to enable downstream analyses.

## Introduction

Hi-C is a genome-wide chromosome conformation capture assay coupled to next-generation sequencing that is used to study 3D genomic organization. Hi-C experiments generate paired-end sequencing data that is used to construct a contact matrix between genomic bins. Intra-chromosomal Hi-C contact maps can be visualized as symmetric heatmaps, with the *x* and *y* coordinates representing genomic locations along the chromosome. Each pixel in the heatmap shows the intensity of chromatin interaction (normalized read count) between the corresponding bins. Examining Hi-C contact map at various resolutions can reveal different scales of 3D organization that are relevant to gene regulation. For example, A/B compartments are usually identified at megabase scale, while topologically associating domains (TADs) are often analyzed using 10-50 kb bins (Szabo et al. [2019]). Significant loop structures can be observed at finer resolutions, using 5-10 kb bins. Mapping fine-resolution contact maps can identify enhancer-promoter interactions that underlie the cell-type-specific regulation of individual genes, and fine-grained contact maps across developmental stages can reveal the dynamics of enhancer rewiring. However, constructing high-resolution contact maps requires high-complexity and deeply sequenced libraries, which can be costly. Contact maps derived from low-coverage libraries often suffer from high noise due to data sparsity, making it challenging to integrate 3D information into the study of gene regulation.

In recent years, the development of single-cell Hi-C technologies has further enabled the study of 3D genome structures at finer cell-type-specific granularity. This advance in principle enables the analysis of 3D structural heterogeneity and variation in regulatory interactions, even across cells of the same cell type. However, analyzing structures at the single-cell or even pseudobulk level remains difficult due to data sparsity. Indeed, it is most common to bin single-cell contact maps at very low resolution such as 100 kb to 1 Mb, or to use of hundreds of thousands of cells to generate pseudobulk data with finer resolutions.

Given the success of deep learning technology for image denoising and image super-resolution (Dong et al. [2014], Tai et al. [2017]), several groups have designed supervised deep learning models to denoise bulk Hi-C libraries. HiCPlus (Zhang et al. [2018]) and HiCNN (Song et al. [2018]) use convolutional neural networks to predict high coverage contact maps from low coverage or downsampled contact maps in the same cell type. Meanwhile, hicGAN (Liu et al. [2019]), DeepHiC (Hong et al. [2020]) and HiCSR (Dimmick et al. [2020]) all use generative adversarial networks (GAN) to impute high resolution data, with DeepHiC and HiCSR employing loss functions specifically tailored to Hi-C data. HiCARN (Hicks and Oluwadare [2022]) uses a cascading neural network to achieve resolution enhancement. Nevertheless, these supervised models often encounter generalization challenges. First, they require paired low-/high-coverage Hi-C libraries for training, limiting their applications in scenarios where suitable training data is not available, such as training to predict at ultra-high resolution where very deeply sequenced libraries are not available, or training on single-cell libraries where high-resolution single-cell contact maps are unknown. Second, they may struggle with generalization to new datasets, for example on datasets where the low-/high-coverage sequencing depth gap between the input data and desired prediction differs from the training scenario. Furthermore, these approaches usually require specific normalization and preprocessing of the Hi-C datasets, posing additional challenges for the post-prediction recovery procedure needed to reconstruct a genome-wide matrix for downstream analysis.

There has been relatively little work on self-supervised denoising strategies, which do not require high-resolution target data during training. Zhang et al. [2022a] introduced DeepLoop, a two-step deep learning framework trained in a self-supervised manner to enhance chromatin interaction detection from low-coverage data. DeepLoop begins with smoothed matrices generated by HiCorr (Lu et al. [2020]), a Hi-C data bias correction pipeline designed to improve interaction detection. The framework involves training two models: LoopDenoise, an autoencoder that is used to denoise the contact matrix; followed by LoopEnhance, a U-Net that uses downsampled HiCorr matrices as input examples and is trained in a supervised manner using enhanced matrices produced by LoopDenoise as targets. While this method improves loop calling from sparse Hi-C matrices, the two-step approach introduces complexity in data preparation and training procedures. The downsampling needed for training LoopEnhance presents a similar challenge to supervised model training, requiring users to train multiple models with different downsampling ratios and to select the one that works best for their data. Moreover, this method has mainly been evaluated for enhancing loop detection but not for denoising other 3D structures like TADs.

For enhancing signals from single-cell Hi-C libraries, Higashi (Zhang et al. [2022b]) and fast-Higashi (Zhang et al. [2022c]) provide fast and efficient algorithms based on hypergraph representation learning, which learns the latent relationship between cells and borrows information from nearby cells to impute cell-type-specific structures. These methods also simultaneously calculate a cell embedding from the imputed 3D structure, providing a convenient option for cell clustering. However, Higashi and fast-Higashi require whole genome 3D information for imputation, posing a memory challenge, with no lower cost option to examine cell-to-cell variability at a local region around a gene of interest.

To address these challenges with a single framework, we introduce HiC2Self, a simple and efficient self-supervised Hi-C denoising model. HiC2Self can be widely used for a variety of tasks, including denoising bulk Hi-C data, denoising Micro-C data at ultra-high resolution, reconstructing cell-cluster-specific contact maps using single-cell Hi-C, and examining local 3D structures at a specific locus or gene at the single-cell level.

## Results

### HiC2Self uses a self-supervised framework to denoise the Hi-C contact maps

An overview of HiC2Self framework is displayed in Figure 1. There are three key components in the HiC2Self framework:

**Figure 1:**
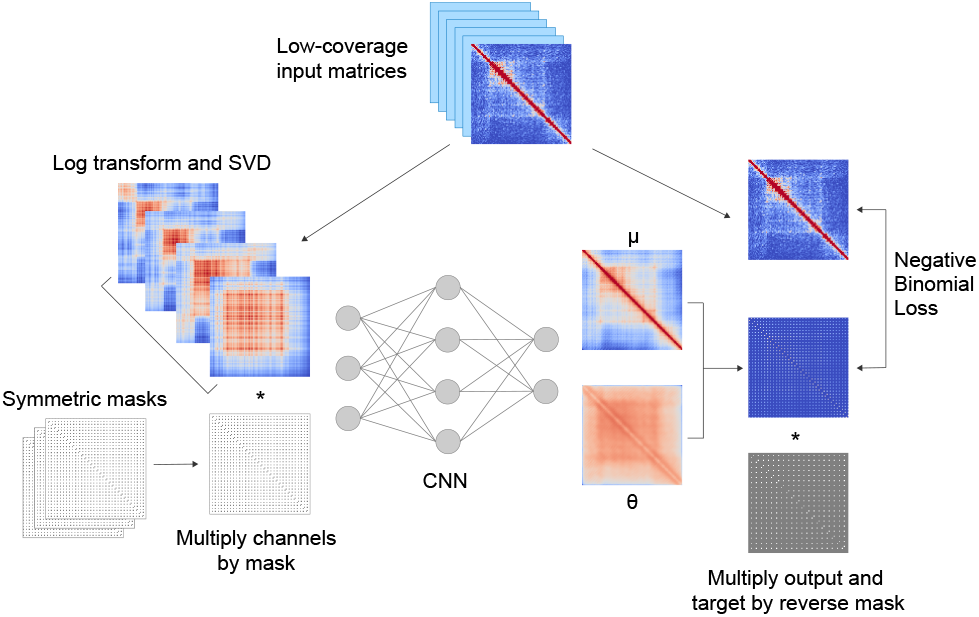
Framework of HiC2Self. HiC2Self uses a self-supervised framework with low-coverage input matrices serving as both the input and the training target; a singular value decomposition on low-coverage inputs to enhance low-rank signals; and symmetric masks to force the model to predict masked pixels from unmasked ones.

- The low-coverage contact matrix is first decomposed and reconstructed using singular value decomposition (SVD) for signal enhancement.
- A self-supervised training framework adapted from Noise2Self (Batson and Royer [2019]) is applied to the data, using the low-coverage input matrices as both the input and the target. The model applies diagonally symmetric grid masks to each input channel and learns to predict the masked pixels from the unmasked pixels.
- The underlying model is a simple convolutional neural network that takes in the multi-channel tensor as the input and has two output channels, representing the mean and dispersion of the negative binomial loss, to directly denoise the contact matrices at the level of raw counts.

As shown in Figure 1, equal-sized low-coverage input matrices are used as both input and training target. During training, each of the low-coverage matrices is log-transformed for numerical stability, then subjected to singular value decomposition (SVD) to obtain the top eigenvalues and eigenvectors. SVD and low-rank reconstruction is a classic approach for 2D image compression and denoising. We reconstructed low-rank approximations of the matrices using the top *n* eigenvalues and their corresponding eigenvectors (for *n* = 1 … 4), with each approximation thus capturing increasing levels of variance in the data.

We adapted the Noise2Self (Batson and Royer [2019]) framework which uses masks to achieve self-supervised training. Masks are matrices of the same dimension as the low-coverage contact maps, where masked regions corresponding to indices in *J* have zero entries, and while unmasked regions at the complementary indices (those in *J*^*C*^) have entries equal to 1. Notationally, we can also identify the set of indices *J* with the corresponding mask matrix, so that the matrix *J*^*C*^ corresponds to the reverse mask (1 − *J*). Since the contact maps are diagonally symmetric, we designed the masks in a similar manner, ensuring that the entries in the masked and unmasked regions are independent from each other. The mask structure is shown in Supplementary Figure S1. Each channel in the multi-channel input matrices is multiplied by the mask before going into the model.

HiC2Self uses a simple convolutional neural network (CNN) to denoise the contact matrices and negative binomial loss to train the model. The CNN model consists of five equal-sized convolutional layers, where each of the first three layers is followed by ReLU activation functions. An exponential function is used at the last layer in order to transform output values back to the raw count space. We assume the raw counts follow a negative binomial distribution. The model outputs two channels, *µ* and *θ*, and we use the negative log-likelihood of negative binomial probability as the training objective to optimize the HiC2Self model. During training, both the predicted channels and the low-coverage input matrix need to be multiplied by the reverse (complementary) mask before calculating the loss (Figure 1).

### HiC2Self accurately reconstructs cell-type-specific contact maps from low-coverage libraries

We first evaluated the denoising performance of HiC2Self on a low-coverage bulk Hi-C library. A relatively shallowly sequenced library on GM12878 (GSE63525, Rao et al. [2014]) with 202.10 million reads was used as the low-coverage input to HiC2Self, and we compared the results against a pooled high-coverage library with 3.5 billion reads that was never seen by the model. Both libraries were binned at 10 kb resolution, and we performed HiC2Self denoising on the low-coverage library on regions up to 1 Mb distance from the diagonal. Two example regions of HiC2Self recovery on chromosome 3 are shown in Figure 2a. Within each example, the top row shows the low-coverage contact maps, followed by the insulation score calculated from the overall contact map. The middle row shows the HiC2Self recovered matrices, and the bottom row shows the unseen pooled high-coverage library. Input and predicted contact maps are KR normalized for visualization. From the visual comparison, HiC2Self recovers structures that are consistent with the high-coverage library. We further examined the identification of topologically associated domains (TADs), called using TopDom (Shin et al. [2016]), on the low-coverage, HiC2Self recovered and high-coverage contact maps. Dashed blue lines show the TAD calls from each contact map. In these examples, the TAD structure in the HiC2Self recovered contact maps is consistent with that from the high-coverage library, while the low-coverage input data can lead to miscalled TADs.

**Figure 2:**
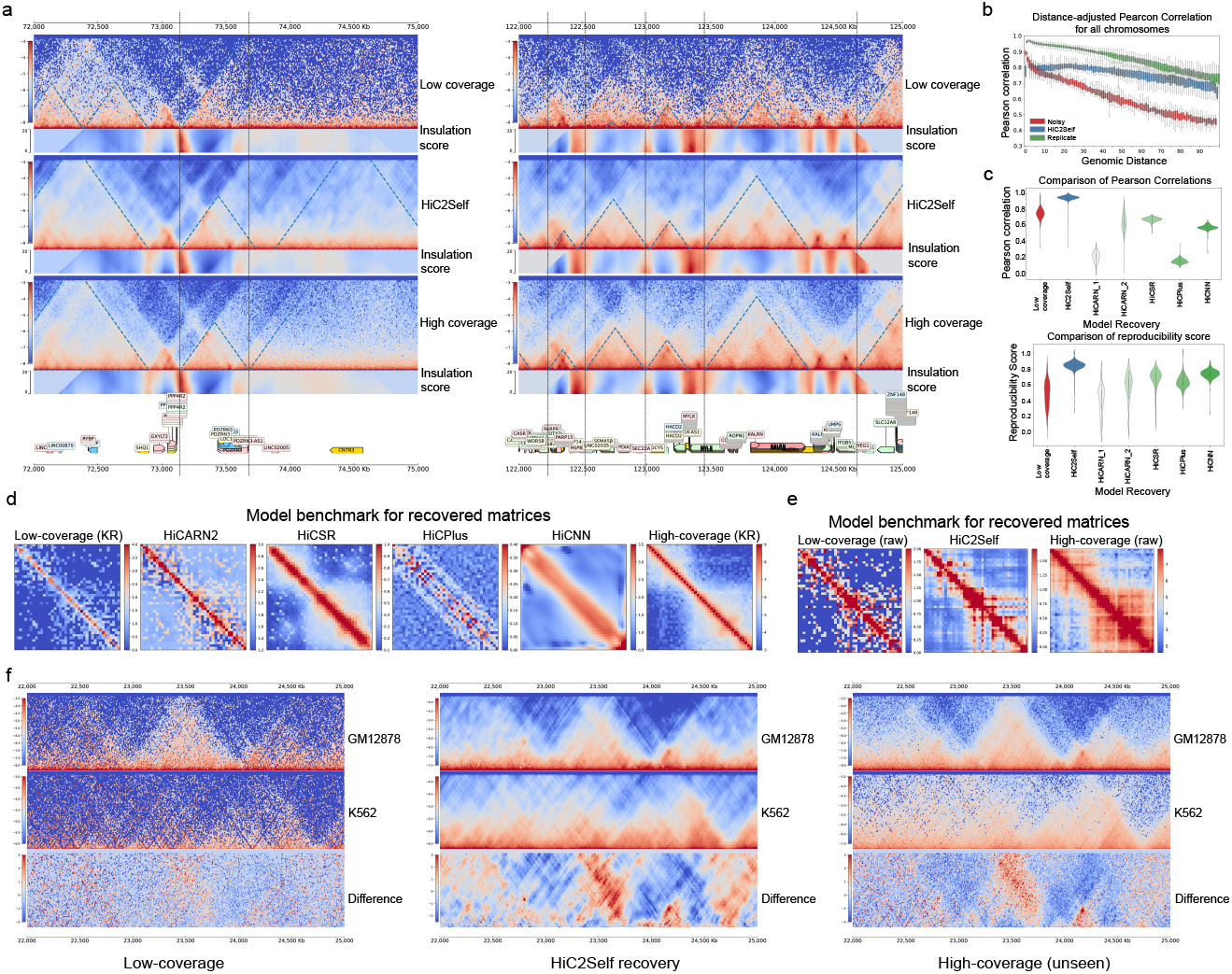
HiC2Self shows robust performance on bulk Hi-C libraries. **(a)** Two random regions on chromosome 3 as a visual comparison example of HiC2Self recovery. The top panel shows the low-coverage Hi-C input, the middle panel shows the HiC2Self recovery, and the bottom shows a deeply sequenced library. The blue dashed triangle lines show the TAD calls identified by TopDom (Shin et al. [2016]), and the vertical dashed lines show the TAD boundaries called from the deeply sequenced library. **(b)** Distance-adjusted Pearson correlation of each library compared with the deeply sequenced library, up to 1 Mb from the diagonal. Red boxes show the correlation from the low-coverage library, blue boxes show the HiC2Self recovery, and green boxes show a deeply sequenced biological replicate compared with the high-coverage library. **(c)** HiC2Self benchmark results against supervised models on an unseen new dataset. The top panel shows the Pearson correlation of each predicted library vs. the deeply sequenced library. The bottom panel shows the reproducibility score (Ursu et al. [2018]). Violin plots from left to right: low-coverage contact map, HiC2Self predicted map, supervised models HiCARN (version 1, Hicks and Oluwadare [2022]), HiCARN (version 2), HiCSR (Dimmick et al. [2020]), HiCPlus (Zhang et al. [2018]), HiCNN (Song et al. [2018]). **(d)** This panel shows an example random region on chromosome 4. From left to right: low-coverage input matrix, recovered matrices from HiCARN version 2, HiCSR, HiCPlus, HiCNN and the corresponding region in the high-coverage contact matrix. All matrices are KR normalized. **(e)** Same example region with HiC2Self. From left to right: low-coverage contact map, HiC2Self recovery, and high-coverage contact map. All matrices are unnormalized. **(f)** A visual comparison of cell-type-specific recovery of HiC2Self. Low-coverage libraries are on the left, HiC2Self recoveries are in the middle, and high-coverage libraries are on the right. In each panel, from top to bottom: GM12878, K562, difference between the two libraries.

For a genome-wide evaluation, we then compared the reconstruction of HiC2Self on this low-coverage dataset against the performance of a deeply sequenced biological replicate library with 2.9 billion reads (GSE63525, Rao et al. [2014]), where again the task was to recover the high-coverage (3.5 billion read) ground truth contact maps. Figure 2b shows a distance-adjusted Pearson correlation across all chromosomes between low-coverage, HiC2Self reconstruction and biological replicate compared with the high-coverage ground truth library. Red bars show the correlation between original low- and high-coverage maps; green bars show the correlation of the biological replicate with the ground truth; and blue bars show the performance of HiC2Self recovered maps. Thus, using a library with less than 0.1% reads as input, HiC2Self was able to achieve almost comparable performance to a deeply sequenced biological replicate library. We also normalized the raw and predicted contact maps of chromosome 3 using HiC-DC+ (Sahin et al. [2021]) and calculated the ROC and PR curves of the significant interactions identified by the low-coverage library, HiC2Self and the biological replicate, compared with the high-coverage unseen library. The results are shown in Supplementary Figure S2a and confirm that HiC2Self can reconstruct significant interactions from a low-coverage library.

We further benchmarked HiC2Self against previously published supervised models for the bulk library denoising task. We compared HiC2Self with four supervised Hi-C contact map denoising models, including HiCSR (Dimmick et al. [2020]), HiCPlus (Zhang et al. [2018]), HiCNN (Song et al. [2018]) and HiCARN (Hicks and Oluwadare [2022]). For supervised models, we downloaded pre-trained checkpoints of models that were trained on low- and high-coverage contact map pairs based on a 1/64 downsampled sequencing depth ratio. The supervised models were trained on GM12878 Hi-C data with KR normalization at 10 kb resolution, with chromosome 4 held out as the test chromosome. We benchmarked all the models on a new low-coverage GM12878 library (84.91 million reads) representing about 1/42 of the sequencing depth of the deeply sequenced library (3.5 billion reads). Chromosome 4 at 10 kb resolution was used for the evaluation of all models. The supervised models were trained and evaluated using KR normalization, whereas HiC2Self was trained with raw matrices and evaluated on matrices with local KR balancing applied to each matrix. We used Pearson correlation and the GenomeDISCO reproducibility score (Ursu et al. [2018]) as evaluation metrics. Figure 2c shows the Pearson correlation and reproducibility score comparing low-coverage input library (red), HiC2Self recovery (blue), and pre-trained HiCARN, HiCSR, HiCPlus, HiCNN recovered matrices (green). HiC2Self outperforms all supervised models when generalizing to a new dataset. For a visual comparison of the denoising results, Figure 2d shows a random example region on the test chromosome (chr4) for all the supervised models (left to right: low-coverage KR normalized matrix, HiCARN2, HiCSR, HiCPlus, HiCNN and high-coverage KR normalized matrix). Figure 2e shows the same region using HiC2Self recovery (left to right: low-coverage raw matrix, HiC2Self recovery, high-coverage raw matrix).

We also compared the significant loop calls from HiC2Self with those from a previous self-supervised training framework called DeepLoop. DeepLoop re-analyzed previously published data from Rao et al. [2014] for mid-range cis-interactions (<2 Mb) at 5 kb resolution. For GM12878, five biological replicates were used for training LoopDenoise, with sequencing depth ranging from 252.43 million reads to 1.78 billion reads. The results are available at (GSE167200). We retrained HiC2Self using only one replicate with 202.10 million reads (GSE63525, Rao et al. [2014]) on chromosome 3 at 5 kb resolution, covering 2 Mb from the diagonal, and calculated significant interactions using HiC-DC+ (Sahin et al. [2021]). The true significant interactions were defined by using HiC-DC+ (Sahin et al. [2021]) on a deeply sequenced library with 3.5 billion reads. DeepLoop achieved an auROC of 0.72 and an auPR of 0.34, whereas HiC2Self obtained an auROC of 0.77 and an auPR of 0.40. Therefore, HiC2Self, trained with a single low-coverage replicate achieved slightly better performance than DeepLoop, which was trained with five replicates with deeper sequencing depth (Supplementary Figure S2b).

As a benefit of its self-supervised training framework, HiC2Self can be easily applied to different datasets and cell types without the generalization challenge seen with supervised models. We again ran HiC2Self on the previously used GM12878 libraries as well as the K562 libraries from Rao et al. [2014], and compared the model performance for capturing cell-type-specific structures. The low-coverage K562 library used as input has 79.91 million reads, while the high-coverage library has 1.4 billion reads. Figure 2f shows a comparison of a cell-type specific region on chromosome 3 of GM12878 (top row in each panel), K562 (middle row), and the difference between the two cell types (bottom row). Low-coverage data are shown on the left, with HiC2Self recovery in the middle, and high-coverage data on the right. The distance-adjusted Pearson and Spearman correlations for chromosome 3 in K562 are shown in Supplementary Figure S2c (left), while the distance-adjusted correlations for cell-type differences (GM12878 - K562) are shown in Supplementary Figure S2c (right). This analysis shows that HiC2Self can accurately reconstruct cell-type-specific structures in different cell types.

### HiC2Self recovers significant interactions at ultra-high resolution

Given the robust performance at accurate recovery of TADs and loops on bulk Hi-C data, we further explored the ability of HiC2Self to reconstruct 3D structures at finer resolution from Micro-C data. Micro-C (Hsieh et al. [2020]) is an assay to map 3D chromosome conformation that uses MNase instead of restriction enzymes to cut the DNA, enabling it to achieve higher resolution compared to Hi-C libraries. We used a low-coverage mouse embryonic stem cell (mESC) Micro-C library with 188.79 million reads, binned at 1 kb resolution, and ran HiC2Self on chromosome 6 to reconstruct the HLA region. A deeply sequenced Micro-C library with 3.4 billion reads was also binned at 1 kb and used as the unseen ground truth to evaluate the reconstruction performance of HiC2Self. Figure 3a shows a visual comparison of the low-coverage library, HiC2Self recovery and the deeply sequenced library. Notably, HiC2Self successfully reconstructed TAD structures from a very sparse low-coverage library. In order to quantitatively evaluate the ability of HiC2Self to reconstruct meaningful structures at 1 kb resolution, we calculated a distance-adjusted z-score for interaction bins at each genomic distance from the diagonal and defined the top 1% of these z-scores as significant interactions in the low-coverage, HiC2Self recovered, and high-coverage contact maps. The ROC and PR curves of interactions identified from the low-coverage library and HiC2Self vs. the high-coverage ground truth are shown in Figure 3b. Thus HiC2Self can reconstruct significant interactions from low-coverage Micro-C data at very fine resolution.

**Figure 3:**
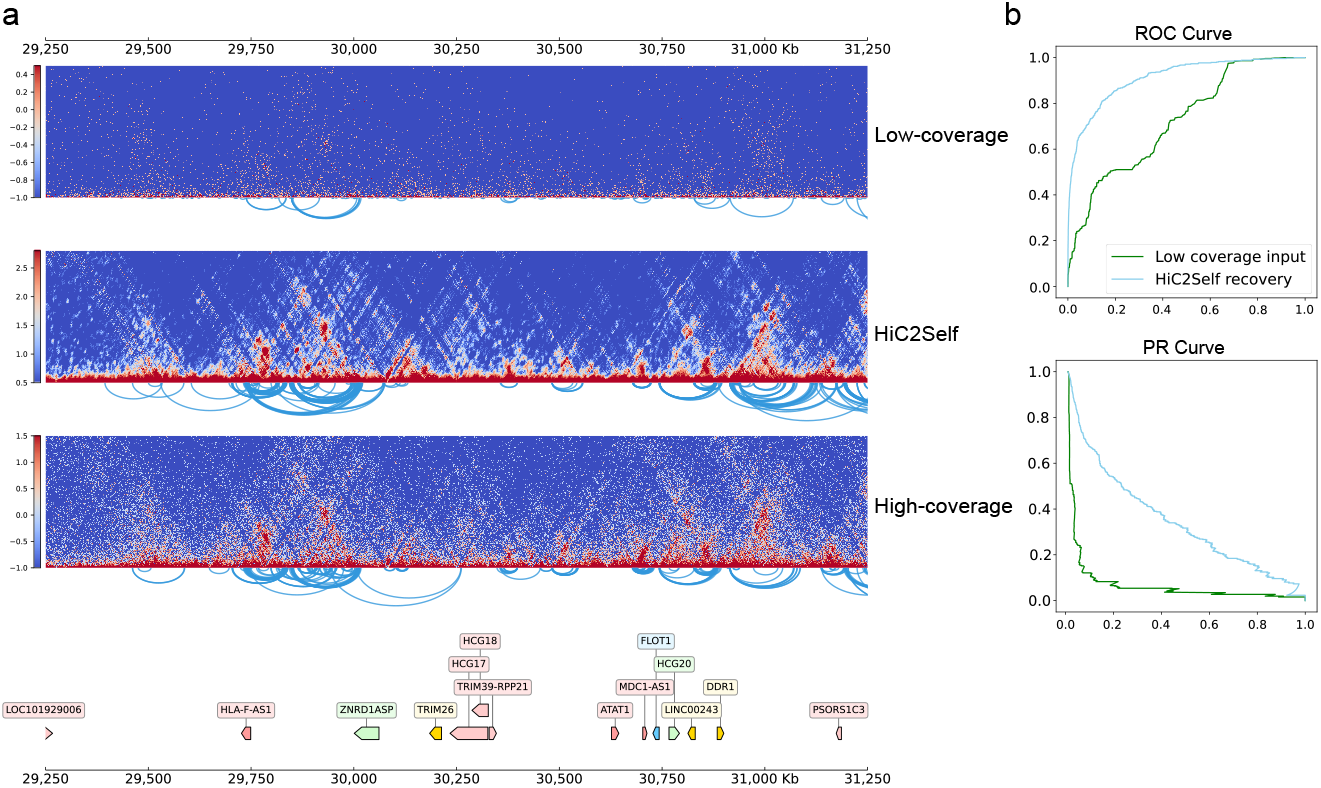
HiC2Self application at ultra-high resolution. **(a)** A visual comparison of the HLA region (chr6: 29,250,000-31,250,000) on chromosome 6 in mouse embryonic stem cells (mESC). Arcs show the significant interactions identified from each data set. The significant interactions are defined as top 1% of the distance-adjusted z-score. **(b)** ROC and PR curve of significant interactions identified from low-coverage data set compared with deeply sequenced data set (green), and HiC2Self vs. deeply sequenced data set (blue).

### HiC2Self accurately reconstructs cell-population specific 3D genome structures with single-cell Hi-C

In addition to achieving robust performance for denoising bulk Hi-C libraries, HiC2Self can also enhance 3D structures from single-cell Hi-C libraries, facilitating the examination of contact maps at the pseudobulk or single-cell level. Single-cell Hi-C technology has progressed rapidly in recent years. In particular, the emergence of various single-cell multi-modal co-assays allows the definition of cell clusters or cell types based on the better understood expression or epigenomic readouts, followed by the examination of cell-cluster-specific pseudobulk 3D structures at high granularity. Single-nucleus methyl-3C sequencing (sn-m3C-seq) (Heffel et al. [2024]) is a single-cell multi-modal assay that simultaneously measures chromosome conformation and DNA methylation and was recently used to profile 53,000 cells from the human pre-frontal cortex (PFC) and hippocampus (HPC) to investigate the epigenome and 3D genome during brain development. In this study, cell types were determined based on clustering and annotation of their methylation profiles.

Using this dataset, we first evaluated the ability of HiC2Self to reconstruct cell-cluster-specific structures around genes of interest based on sparse pseudobulk input data over a few dozen cells. We examined a 4 Mb local region around *RORB*, an important gene related to schizophrenia, during early human brain development. We randomly sampled 50 cells from each cell subtype in the prefrontal cortex (PFC), including radial glia cells in the second trimester (2T-RG-1) and medial ganglionic eminence (eMGE) at the prenatal stage, and layer 1-3 cortical neurons expressing *NRXN2* (L1-3-NRXN2), layer 4-5 neurons expressing *FOXP2* (L4-5-FOXP2), oligodendrocyte cells (ODC) at the adult stage, and binned the 50-cell pseudobulk contact maps at 10 kb resolution. We then trained HiC2Self on chromosome 9 of this 50-cell pseudobulk input contact map. Figure 4a shows a visual comparison of the 4 Mb region around the *RORB* gene, with the raw 50-cell pseudobulk maps on the left, HiC2Self recovered maps in the middle, and 25,000-cell pseudobulk maps for each cell type as the ground truth on the right. A similarity score comparison of the raw 50-cell pseudobulk map and HiC2Self recovery vs. the 25,000-cell pseudobulk map is shown in Figure 4b. Furthermore, zooming into a 1 Mb region around the *RORB* gene, it is clear that HiC2Self successfully reconstructed the distinct structure at the L4-5-FOXP2 cell stage with as little as 50 cells. Thus, HiC2Self applied to pseudobulk maps of a few dozen cells enables the examination of 3D genome folding structures and regulatory interactions around genes of interest in rare cell types from single-cell Hi-C datasets.

**Figure 4:**
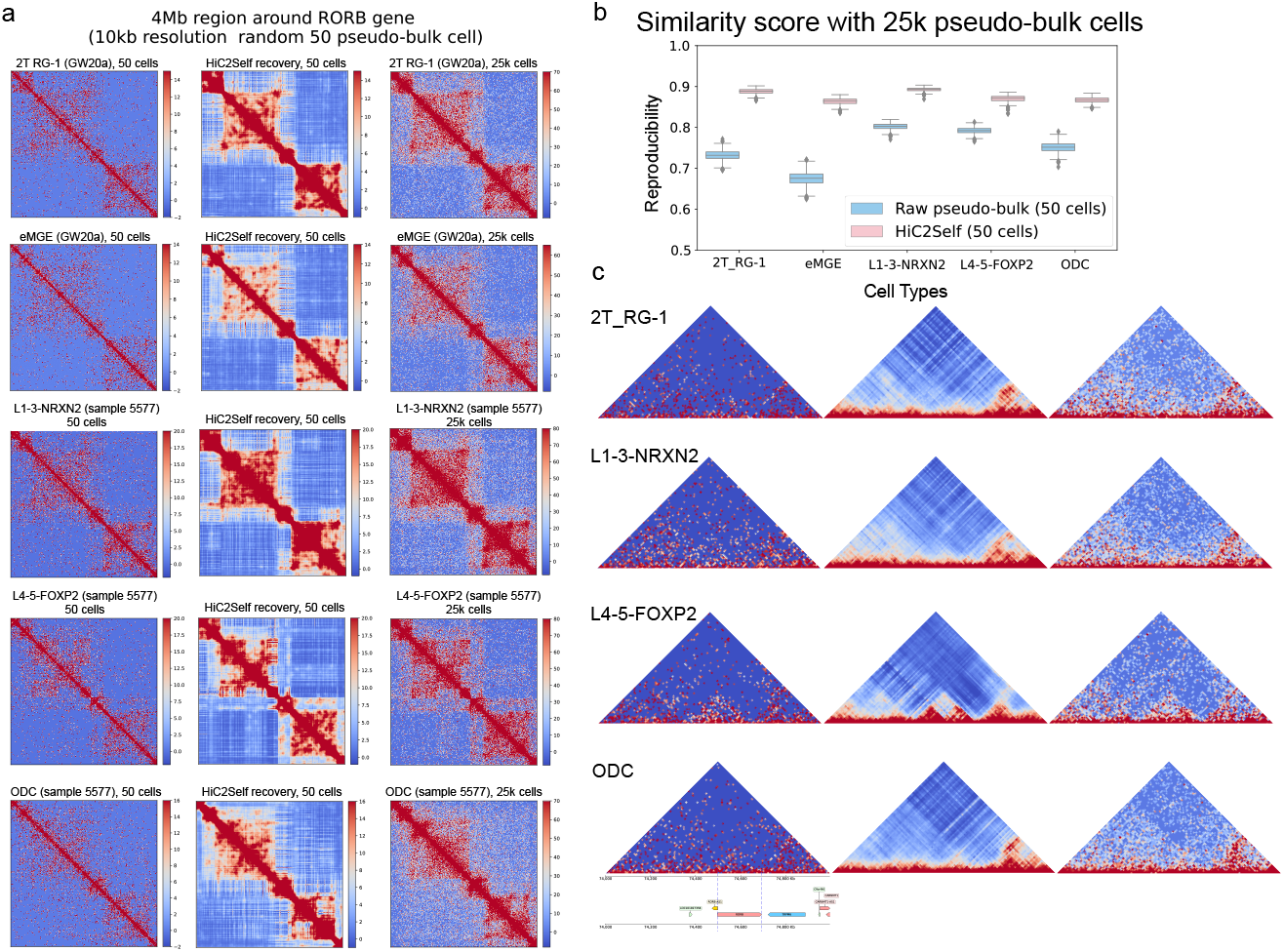
HiC2Self recovers cell-population-specific 3D structures from single-cell Hi-C libraries. **(a)** A comparison of a 4 Mb region around the *RORB* gene, with pseudo-bulk libraries binned at 10 kb resolution. The left column shows the pseudobulk map based on 50 cells, the middle column shows the HiC2Self recovery, and the right column shows the higher coverage pseudobulk map based on 25,000 cells. Different cell types are shown from top to bottom: 2T_RG-1 (radial glia cells in the second trimester), eMGE_GW20a (medial ganglionic eminence during gestational week 20), L1-3-NRXN2 (layer 1-3 cortical neurons expressing *NRXN2*), L4-5-FOXP2 (layer 4-5 neurons expressing *FOXP2*), ODC_5577 (oligodendrocyte cells). **(b)** The similarity score comparing 50-cell pseudobulk maps vs. the 25,000-cell pseudobulk maps (blue), and the HiC2Self recovered 50-cell maps vs. the 25,000-cell pseudobulk maps (pink). **(c)** A zoomed-in visualization of a 1 Mb region around the *RORB* gene, where each row shows the cell-type-specific 50-cell pseudo-bulk, HiC2Self recovery, and 25,000-cell pseudobulk structures. The gene annotation track is shown at the bottom.

### HiC2Self reconstructs single-cell 3D structure without borrowing information from neighboring cells

Given the strong performance of HiC2Self in recovering structure from low cell number pseudobulk maps in single-cell Hi-C datasets, we tested the application of HiC2Self at the single-cell level. In the human cortex sn-m3C-seq dataset (Heffel et al. [2024]), we noticed that some genes (e.g. *DLG2* gene) show strong variability in methylation levels even within the same cell type. Therefore, in addition to reconstructing the cell-type-specific 3D structures, we investigated whether HiC2Self couldcapture cell-to-cell variation in the local 3D structure around genes of interest in individual cells of the same cell type. Figure 5a shows a 20 Mb region around the *DLG2* gene using a pseudobulk map of 1,248 Exc-CA (excitatory neurons from the Cornu Ammonis regions) cells from the hippocampus (HPC) region, binned at 50 kb resolution. Figure 5b shows the methylation level of *DLG2* and nearby genes from the top 5% highest *DLG2*-methylated cells versus the bottom 5% *DLG2*-methylated cells. We can see that even within the same cell type, cells present diverse methylation patterns for certain genes. Figure 5c shows the 3D structure of a 20 Mb region around the *DLG2* gene from 5% lowest methylated cells (top), the middle 5% of cells with intermediate methylation (47.5-52.5%, middle) and 5% highest methylated cells (bottom). Cells with higher methylation at *DLG2* tend to form longer range interactions compared with cells with low methylation at this gene. Figure 5d shows two random cells sampled from the 5% lowest *DLG2*-methylated cells, with the sparse raw single-cell contact map on the left, Gaussian smoothed map in the middle, and HiC2Self reconstructed single-cell map on the right. Figure 5e shows two random cells sampled from 5% highest *DLG2*-methylated cells. We further calculated a UMAP using the raw single-cell (top) and the HiC2Self reconstructed (bottom) local contact maps, shown left to right in Supplementary Figure S3: the UMAP colored by developmental stage, contact scores reported from Heffel et al. [2024] and mCG and mCH methylation of the *DLG2* gene. The UMAP shows that cells grouped by the local 3D structure, as recovered by HiC2Self around the *DLG2* gene, show a pattern consistent with its methylation levels. HiC2Self thus revealed cell-to-cell variability in 3D structure within the same cell type that correlated with the methylation pattern, making possible the study of single-cell gene regulation patterns among cells of the same cell type.

**Figure 5:**
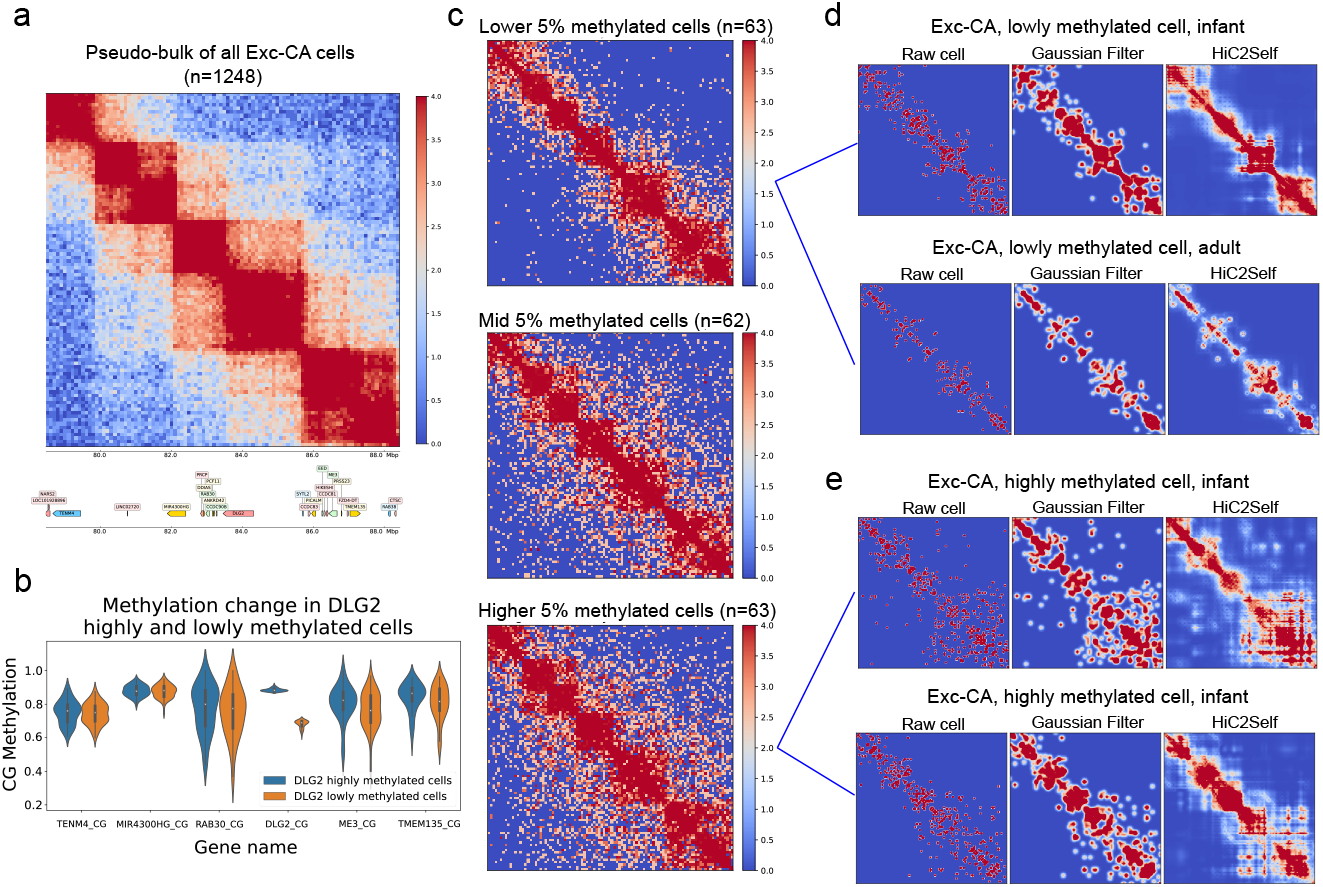
HiC2Self reconstructs the single-cell 3D structures. **(a)** A 20 Mb region around the *DLG2* gene from a pseudobulk contact map of 1,248 Exc-CA cells binned at 50 kb resolution. The gene annotation track is shown at the bottom. **(b)** The mCG methylation level of *DLG2* and nearby genes (including *TENM4, MIR4300HG, RAB30, PDLG2, ME3* and *TMEM135*) from the lowest 10% *DLG2*-methylated cells (orange) vs. the highest 10% *DLG2*-methylated cells (blue). **(c)** Comparison of the 20Mb region around the *DLG2* gene from the pseudobulk contact map with lower 5% DLG2-methylated cells (top), middle 5% *DLG2*-methylated cells (middle), and highest 5% *DLG2*-methylated cells (bottom). Each pseudobulk library consists of 62-63 cells. **(d)** Two randomly sampled cells from the lowest 5% *DLG2*-methylated cells, with raw cell contact matrix on the left, Gaussian smoothed matrix in the middle, and HiC2Self recovered matrix on the right. **(e)** Two randomly sampled cells from the highest 5% *DLG2*-methylated cells, with raw cell matrix on the left, Gaussian smoothed matrix in the middle, and HiC2Self recovery on the right.

Finally, we benchmarked HiC2Self with a previously published model for scHi-C imputation called Higashi (Zhang et al. [2022b]), evaluating on chromosome 1 of the sn-m3C-seq PFC (prefrontal cortex) dataset (Heffel et al. [2024]) at 500 kb resolution. We trained HiC2Self with only one chromosome, while Higashi incorporates all the chromosomes during training. Supplementary Figure S4a shows two random example cells from the Exc_L1-3 (excitatory neurons localized in cortical layers 1-3) cell type at 500 kb resolution, with (left to right) as raw single-cell contact map, HiC2Self recovery, Higashi recovery, and pseudobulk of all the cells from this cell type. We further used distance-adjusted correlation and the reproducibility score to evaluate the performance of two models, with results shown in Supplementary Figure S4b. In this comparison, we selected three cell types with distinct 3D structures, namely Exc_L1-3, Inh_MGE (inhibitory neurons originating from the medial ganglionic eminence (MGE)), NonN_Astro (Non-Neuronal Astrocytes), and evaluated all the cells from each of these three cell types against the pseudo-bulk of all three cell types to see whether the models could reconstruct cell-type-specific structures, and whether reconstructed single-cell maps were most similar to the pseudobulk map of the corresponding cell type. The top row shows the averaged distance-adjusted Spearman correlation, which is calculated for each single cell by first calculating the correlation at each genomic distance from the diagonal, then averaging them to obtain an overall distance-adjusted Spearman correlation for that cell. The bottom row shows the reproducibility score. From left to right are the performance of raw single-cell contact maps, Gaussian smoothed maps, HiC2Self recovery and Higashi recovery. In each panel, boxes show the correlation or similarity of all the cells from Exc_L1-3, Inh_MGE, NonN_Astro vs. pseudobulk of Exc_L1-3, and all the cells from Exc_L1-3, Inh_MGE, NonN_Astro vs. pseudo-bulk of Inh_MGE, etc. Using distance-adjusted Spearman correlation, all the cell types showed strong cell-type-specific structures, where HiC2Self-recovered single-cell maps had the strongest correlation with the pseudobulk of the corresponding cell type. Higashi had the highest Spearman correlation compared with other methods. Using the reproducibility score, however, HiC2Self slightly outperformed Higashi in similarity to the correct pseudobulk cell type (p=3.55 × 10^*−*161^, Wilcoxon one-sided ranksum test), as well as giving superior cell-type-specificity (i.e., higher similarity to the corresponding pseudobulk map than those of other cell types).

## Discussion

In this study, we present HiC2Self, a simple and efficient self-supervised method designed to denoise bulk and single-cell Hi-C contact maps to empower downstream analyses. The HiC2Self framework includes three key components: a) a convolutional neural network with SVD reconstructed channels for structural enhancement of input contact maps; b) a self-supervised framework involving masks to use low-coverage data as both the input and training target; and c) a negative-binomial loss to run the model with raw count matrices. In the Results section, we have shown that HiC2Self can accomplish various tasks, including denoising low-coverage bulk Hi-C data, recovering significant interactions at ultra-high resolution from Micro-C data, reconstructing cell-population-specific 3D genome structure in low cell number pseudobulk maps from single-cell Hi-C, and examining gene regulatory patterns at genes of interest at the single-cell level.

In addition to using the negative log-likelihood (NLL) of negative binomial probability as the training loss as discussed earlier, we also experimented with different loss functions. We first replaced the negative binomial loss with the structural similarity index (SSIM, Wang et al. [2004]) loss instead. SSIM is a perceptual metric to measure the similarity between two images, which is commonly used in the computer vision domain. This loss prioritizes the maintenance of structural similarity instead of penalizing equally for pixel-wise differences. In particular, we applied the SSIM loss on our single-cell datasets and found a notable capacity to recover intricate global structures, as compared with primarily local pattern recovery using traditional NLL loss. However, while the enhanced structures looked promising, they appeared exaggerated in regions with high sparsity, where potentially there was insufficient information to correctly recover the true structures. We also applied SSIM loss to bulk Hi-C contact maps and compared with the unseen deeply sequenced ground truth maps and found the performance was not as good as the NLL loss, potentially indicating the need for more hyper-parameter tuning. Therefore, we retained the NLL of negative-binomial loss function for all denoising tasks.

Inspired by the SSIM loss design, we further experimented with training the model in a multi-resolution manner to capture both local and global structures. In this setting, we not only optimized the loss function on the original resolution to capture local structures but also performed max-pooling on the prediction and target matrices, then optimized the loss on the lower dimensional matrices to capture global structures. We applied this design to both bulk and single-cell applications and found that its performance varied across datasets but was overall comparable to single-resolution training. As a result, we opted to use single-resolution training as the default, while providing multi-resolution training as an optional feature in the codebase.

While HiC2Self is capable of addressing numerous applications, it also displays certain limitations. First, due to the design of the SVD reconstructed channel and masks, HiC2Self requires diagonally symmetric contact maps as input. This design may increase the computational burden when we want to recover long-distance structures, as a bigger input matrix is required to train the model. This design also brings difficulties with denoising trans-contacts, where symmetric matrices are not available, although an analogous self-supervised strategy may be feasible. Second, although HiC2Self shows strong performance for reconstructing structures from sparse contact maps, it cannot impute structures that are missing due to limitations of the sequencing assay. For example, when reconstructing 3D structures at 1 kb resolution, HiC2Self cannot impute the unmappable regions from Hi-C libraries. Further model development to address these challenges would be of interest.

## Conclusion

We have presented HiC2Self, an efficient self-supervised deep learning framework designed to denoise Hi-C contact maps. HiC2Self’s desnoising strategy includes three key design decisions: a convolutional model with SVD reconstructed channels, a self-supervised training framework achieved by masks, and negative-binomial training loss to train on raw count matrices. HiC2Self can be widely used in a variety of tasks, including denoising bulk Hi-C datasets, reconstructing biologically meaningful 3D structures from Micro-C data at high resolution, as well as reconstructing pseudobulk or single-cell 3D structures to study cell-to-cell variability in scHi-C data sets. HiC2Self thus provides a flexible denoising tool to enhance the interpretation of single-cell and bulk 3D genomics datasets.

## Methods

### Data Availability

#### Bulk GM12878 dataset

High coverage Hi-C datasets are generated by sequencing multiple libraries and aggregating read counts across libraries. To obtain low-coverage Hi-C training data, we generated a contact map from a single library of a high-coverage Hi-C dataset and evaluated performance against the aggregated multi-library map. Intra-chromosomal Hi-C raw count contact maps were generated without normalization. For each chromosome in the low-coverage dataset, we further extracted equal-sized square submatrices along the diagonal, representing genomic interactions up to 1 Mb in linear distance. These symmetric submatrices *X* were used as the training set for our model. HiC2Self was trained and evaluated on low- and high-coverage Hi-C data, respectively, as described above. Low-/high-coverage raw count matrices for the ENCODE GM12878 cell line were downloaded from GEO (GSE63525, Rao et al. [2014]). A single low-coverage library (experiment HIC001) with 202.10 million reads was used as low-coverage data to train the model, and pooled primary libraries with 3.5B reads (low/high ratio = 1*/*18) were used as high-coverage Hi-C data to evaluate model performance. Raw count data were downloaded in .hic format and further binned at 10 kb resolution matrix using Juicer (Durand et al. [2016]). Libraries were aligned to hg19. Equal-sized (200 × 200) submatrices were extracted along the diagonal from intra-chromosomal low-coverage Hi-C contact maps to train the model.

#### Bulk K562 dataset

Low-/high-coverage raw count matrices for the K562 cell line were downloaded from GEO (GSE63525, Rao et al. [2014]). The low-coverage library (experiment HIC071) with 79.9M reads was used as low-coverage input for model training, while the pooled high-coverage library with 1.4B reads (low/high ratio=1/17) was used to evaluate model performance. Both libraries were aligned to hg19, and the matrices were processed in the same way as the bulk GM12878 data described above.

#### DeepLoop analysis results on GM12878

DeepLoop (Zhang et al. [2022a]) re-analysis of previously published Hi-C libraries (GSE63525, Rao et al. [2014]) can be found on GSE167200. We downloaded the file GSE167200_GM12878.in_situ.raw_HiCorr_LoopDenoise.txt.gz for the reported loop calls on the GM12878 in situ library.

#### Bulk Micro-C library for ultra-high resolution denoising

Both the low- and high-coverage libraries are downloaded from the 4DN data portal (Reiff et al. [2022]). A high-coverage library on H1 derived differentiated endoderm was downloaded from 4DNESW1SPPTD and contained 3.4 billion reads, while a low-coverage Micro-C library was downloaded from 4DNESP4MARXG and contained 188.79 million reads. These libraries were aligned to mm10. The contact maps were unnormalized and binned at 1 kb resolution, where equal-sized square matrices along the diagonal were extracted from chromosome 6. Each matrix had a shape of 400∗ 400, which covered 400 kb from the diagonal. In the evaluation of significant interactions, we calculated a distance-adjusted z-score (observed counts - average counts for that distance)/(standard deviation of counts for that distance) and defined significant interactions as the top 1% of the interactions from each library.

#### Single-nucleus methyl-3C sequencing (sn-m3C-seq) dataset

A sn-m3C-seq dataset of human brain development was downloaded from GSE213950 with hg38 alignment. The human sample and metadata for cell types were matched from the Supplementary Table 1 and 2 from the original publication Heffel et al. [2024]. For the HPC data, we downloaded specimen UA19-27, UA18-22, GW18-Ant, GW18-Post, GW20 and UMB4267. We randomly sampled 50 cells from each cell type and extracted 400 × 400 sized matrices around the *RORB* gene on chromosome 9 as training data. For the PFC data, we downloaded specimen UMB1863, UMB5577 and GW20a, and only Exc-CA cells were extracted for the study.

#### sn-m3C-seq PFC dataset

The sn-m3C-seq PFC dataset (Heffel et al. [2024]) used to benchmark with Higashi was downloaded from the pre-processed data in [Higashi tutorial](https://github.com/ma-compbio/Fast-Higashi/blob/main/PFC%20tutorial.ipynb). We converted the pairs file into contact matrices and extracted 400 × 400 sized matrices along the diagonal to train HiC2Self.

#### Self-supervised framework

Noise2Self (Batson and Royer [2019]) is a self-supervised denoising framework that uses 𝒥 -invariant functions *f*, where 𝒥 represents the partition of the input data dimensions *m* into subsets, and we consider a subset *J* ∈ 𝒥 and its complement *J*^*C*^. Given an unseen clean signal *y* ∈ ℝ^*m*^, we assume that *x* is a mean-zero noisy observation, where 𝔼 [*x*|*y*] = *y*. For any fixed subset *J*, we further assume that a noisy observation on subspace *x*_*J*_ is independent of the one on its complement 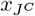 given *y*. With these two assumptions, a function *f* : ℝ^*m*^ → ℝ^*m*^ is defined as 𝒥-invariant if *f* (*x*)_*J*_ is independent of *x*_*J*_ for every *J* ∈ 𝒥.

The ordinary denoising loss function is defined as

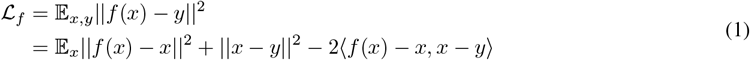

which is the sum of a self-supervised loss and the variance of the noise. With a *J* -invariant function *f* and the previous assumptions, this simplifies to

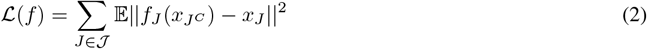

so that the denoising function *f* can be optimized using only noisy observations *x*.

The 𝒥 -invariance property is realized using masks. We denote the masked area as *x*_*J*_ and the unmasked area as 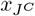. Given the symmetric nature of Hi-C contact maps and the requirement that 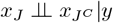, we designed masks that are symmetric with respect to the diagonal.

#### Model architecture

HiC2Self uses a simple convolutional neural network (CNN), as shown in Figure 1. Within the model, raw count input matrices *X* were first log2-transformed 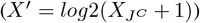 in order to guarantee numerical stability for subsequent steps.

Singular value decomposition (SVD) and low-rank reconstruction is a classic approach for 2D image compression and denoising. In order to enhance the low-rank structures extracted from low-coverage submatrices in the log2-transformed space, we performed SVD on the log2-transformed matrices *X*^*′*^ = *U* Σ*U*^*T*^, generated reconstructions 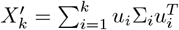 using the top *k* eigenvectors, *k* ∈ [1, 4], and concatenated these matrices with *X*^*′*^ as additional input channels for the CNN.

The convolutional part of the model consists of five equal-sized convolutional layers, where each of the first three layers is followed by ReLU activation functions. An exponential function was used as the activation function for layer 4 and 5 in order to transform output values back into raw count space.

#### Loss function

##### Negative log-likelihood of negative binomial loss

Inspired by the deep count autoencoder (DCA) model for single-cell data (Eraslan et al. [2019]), we used a negative binomial loss for the raw count matrices to train our model. We assume that the count from each bin (*x*_*ij*_) of the contact map *X* follows a negative binomial distribution with parameters *µ*_*ij*_ and *θ*_*ij*_, *x*_*ij*_ ∼ *NB*(*µ*_*ij*_, *θ*_*ij*_). The loss function is defined as

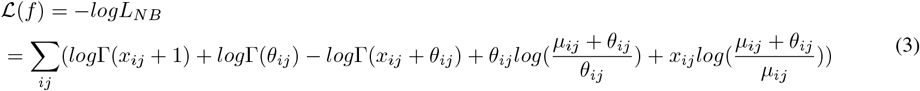

As shown in Figure 1, HiC2Self outputs two channels, corresponding to *µ* and *θ* in the loss function above. We use *µ*_*ij*_, the expected value for each bin *x*_*ij*_, as the predicted value for our denoising results.

##### Genome-wide prediction

HiC2Self produces denoised results as raw counts for fixed-sized submatrices, which can easily be assembled into a whole-chromosome prediction. To do this, we extracted submatrices along the diagonal, consecutively striding by one bin each time. Denoised results were generated for each submatrix, and predicted counts for overlapping submatrices were averaged. The resulting predicted high coverage results were saved as a .hic file using Juicer tools (Durand et al. [2016]), or as a .cool file using [HiCPeaks python implementation](https://github.com/XiaoTaoWang/HiCPeaks) for downstream analysis.

### Model availability

Data preparation pipeline and model scripts are available at github.com/ruy204/HiC2Self.

## Supplementary Figures

**Figure S1:**
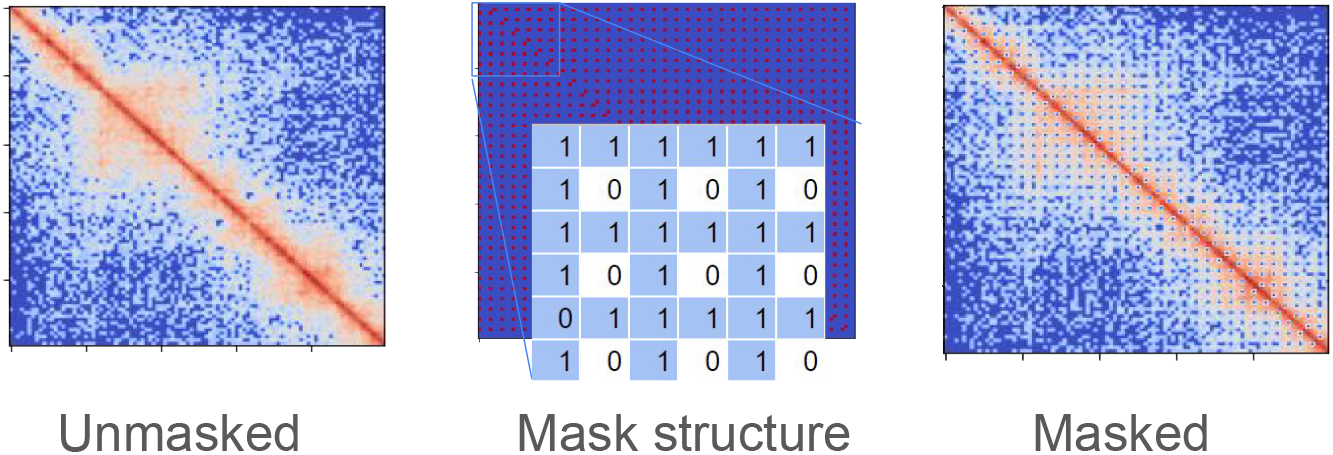
Mask structure of HiC2Self. (Left) Example low-coverage input matrix without a mask. (Middle) Diagram and a zoomed-in view of the mask structure. The mask is diagonally-symmetric, where 0 entries are in the masked regions and 1 entries are in the unmasked regions. (Right) Same example input matrix multiplied by the mask.

**Figure S2:**
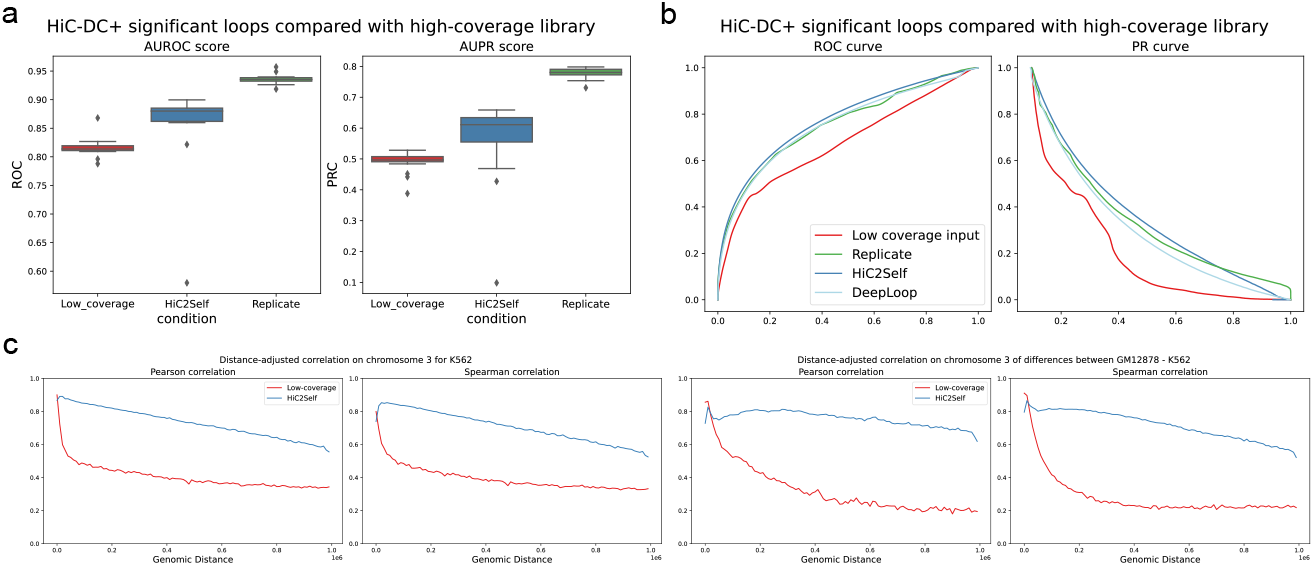
Evaluation of HiC2Self on bulk Hi-C data. **a** AUROC and AUPR of significant interactions. All libraries are bulk GM12878 binned at 10 kb resolution, and significant interactions are identified using HiC-DC+ (Sahin et al. [2021]). Red boxes show the AUROC and AUPR score of significant interactions comparing low-coverage vs. high-coverage libraries, across all chromosomes. Blue boxes show the performance of HiC2Self, and green boxes show the results of a deeply sequenced biological replicate Hi-C library. **b** ROC and PR curves for mid-range (<=2 Mb) cis-interactions at 5 kb resolution, comparing DeepLoop and HiC2Self. Significant interactions are identified by HiC-DC+ (Sahin et al. [2021]). The plot shows the performance of low-coverage data (red line, auROC=0.67, auPR=0.26), a deeply sequenced biological replicate (green line, auROC=0.75, auPR=0.39), DeepLoop trained with five replicates with sequencing depth ranging from 252.43 million to 1.78 billion reads (light blue, auROC=0.72, auPR=0.34), and HiC2Self trained with one low-coverage replicate with a sequencing depth of 202.10 million reads (dark blue, auROC=0.77, auPR=0.40). **c** Distance-adjusted correlations for cell-type specificity. (Left) Distance-adjusted Pearson and Spearman correlation for chromosome 3 of K562, showing the correlation between low-coverage data vs. high-coverage data (red line) and HiC2Self recovery vs. high-coverage data (blue line). (Right) Distance-adjusted Pearson and Spearman correlation of the structural differences between GM12878 and K562. The difference is calculated by GM12878 - K562.

**Figure S3:**
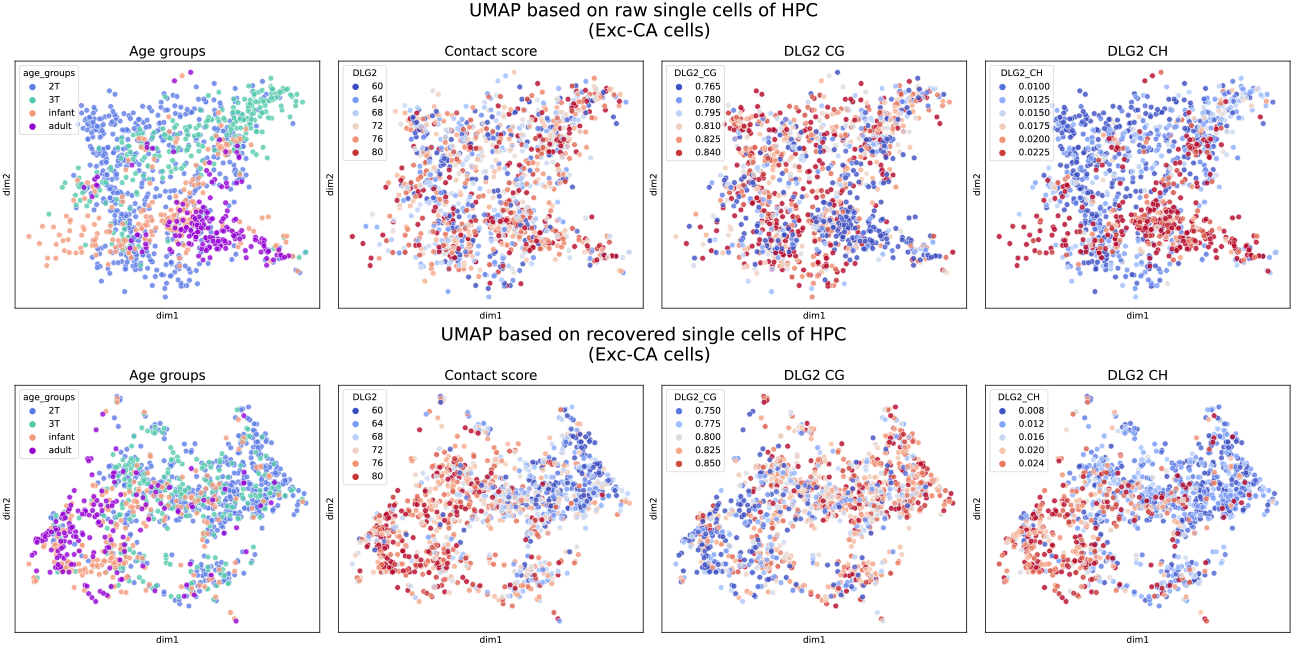
UMAP clustering with local 3D structure of 20 Mb around DLG2 gene. All the cells are from the same cell type, Exc-CA. (Top) UMAP using the local 3D structure within 20 Mb around the *DLG2* gene from raw single cells. (Bottom) UMAP using the 20 Mb region local structure from HiC2Self recovered single-cell contact maps.

**Figure S4:**
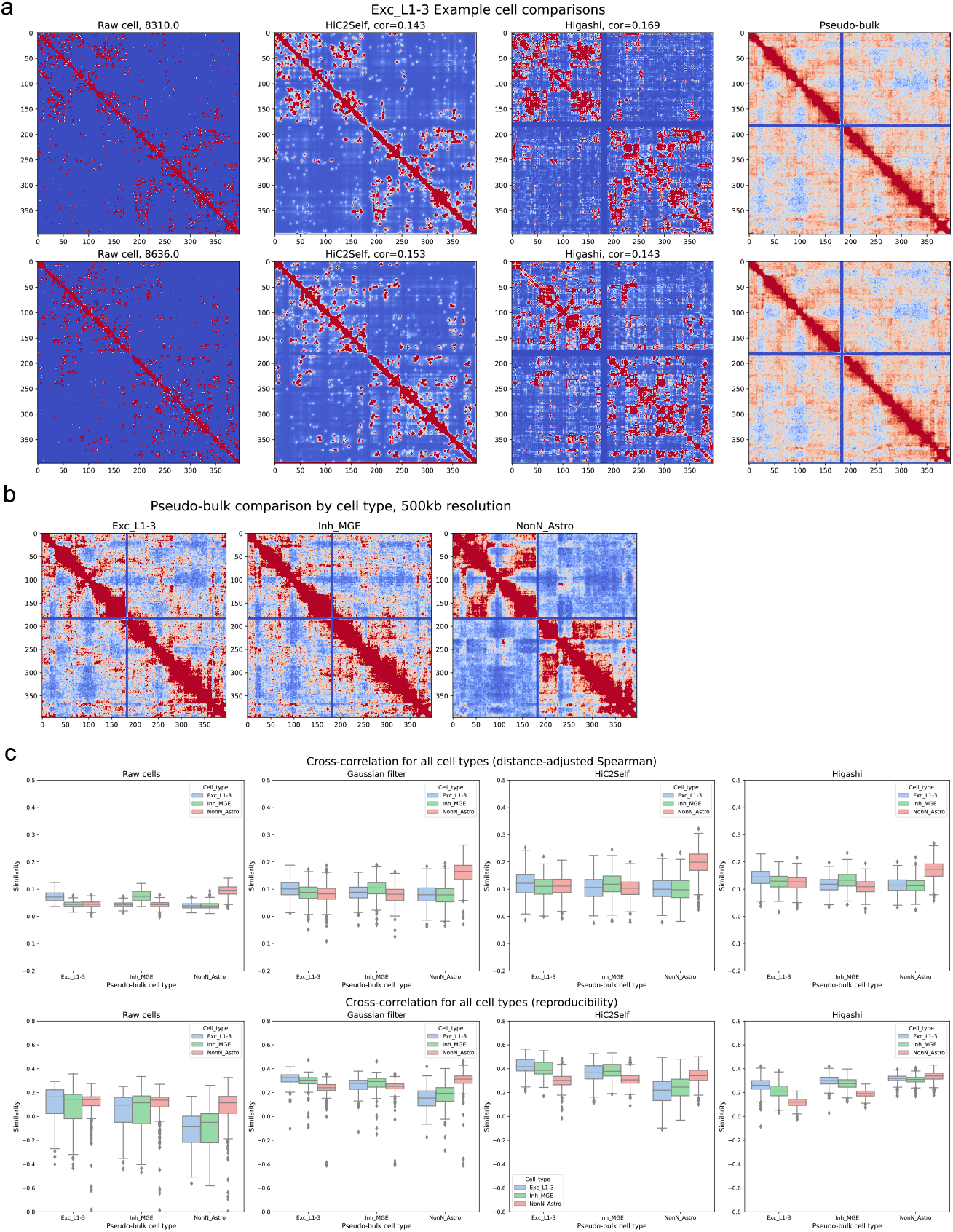
Benchmark with Higashi at 500 kb resolution. **(a)** Two random example cells from the Exc_L1-3 cell type, showing left to right: raw single-cell contact map, HiC2Self recovery, Higashi recovery, pseudobulk contact map for this cell type. The top row shows an example cell where the distance-adjusted Spearman correlation is higher in Higashi, and the bottom row shows an example cell where the correlation is higher is HiC2Self. **(b)** A visual comparison of the pseudobulk cells from three distinct cell types (Exc_L1-3, Inh_MGE and NonN_Astro). **(c)** Cross-comparison of cells from these three distinct cell types (Exc_L1-3, Inh_MGE and NonN_Astro) compared with the corresponding pseudobulk map. Left to right: comparison of raw single-cell maps, Gaussian smoothed cells, HiC2Self recovery and Higashi recovery. The top row shows the distance-adjusted Spearman correlation compared with the pseudo-bulk, and the bottom row shows the reproducibility score.

## References

Quentin Szabo, Frédéric Bantignies, and Giacomo Cavalli. Principles of genome folding into topologically associating domains. Science Advances, 5(4):eaaw1668, April 2019. ISSN 2375-2548. doi:10.1126/sciadv.aaw1668. URL https://www.science.org/doi/10.1126/sciadv.aaw1668.

Chao Dong, Chen Change Loy, Kaiming He, and Xiaoou Tang. Learning a deep convolutional network for image super-resolution. In David Fleet, Tomas Pajdla, Bernt Schiele, and Tinne Tuytelaars, editors, Computer Vision – ECCV 2014, pages 184–199, Cham, 2014. Springer International Publishing. ISBN 978-3-319-10593-2.

Ying Tai, Jian Yang, and Xiaoming Liu. Image super-resolution via deep recursive residual network. In 2017 IEEE Conference on Computer Vision and Pattern Recognition (CVPR), pages 2790–2798, 2017. doi:10.1109/CVPR.2017.298.

Yan Zhang, Lin An, Jie Xu, Bo Zhang, W. Jim Zheng, Ming Hu, Jijun Tang, and Feng Yue. Enhancing Hi-C data resolution with deep convolutional neural network HiCPlus. Nature Communications, 9(1):750, February 2018. ISSN 2041-1723. doi:10.1038/s41467-018-03113-2. URL https://www.nature.com/articles/s41467-018-03113-2.

Tzu-An Song, Samadrita Roy Chowdhury, Kyungsang Kim, Kuang Gong, Georges El Fakhri, Quanzheng Li, and Joyita Dutta. Super-resolution pet using a very deep convolutional neural network. In 2018 IEEE Nuclear Science Symposium and Medical Imaging Conference Proceedings (NSS/MIC), pages 1–2, 2018. doi:10.1109/NSSMIC.2018.8824683.

Qiao Liu, Hairong Lv, and Rui Jiang. hicGAN infers super resolution Hi-C data with generative adversarial networks. Bioinformatics, 35(14):i99–i107, 07 2019. ISSN 1367-4803. doi:10.1093/bioinformatics/btz317. URL https://doi.org/10.1093/bioinformatics/btz317.

Hao Hong, Shuai Jiang, Hao Li, Guifang Du, Yu Sun, Huan Tao, Cheng Quan, Chenghui Zhao, Ruijiang Li, Wanying Li, Xiaoyao Yin, Yangchen Huang, Cheng Li, Hebing Chen, and Xiaochen Bo. Deephic: A generative adversarial network for enhancing hi-c data resolution. PLOS Computational Biology, 16(2):1–25, 02 2020. doi:10.1371/journal.pcbi.1007287. URL https://doi.org/10.1371/journal.pcbi.1007287.

Michael C. Dimmick, Leo J. Lee, and Brendan J. Frey. Hicsr: a hi-c super-resolution framework for producing highly realistic contact maps. bioRxiv, 2020. doi:10.1101/2020.02.24.961714. URL https://www.biorxiv.org/content/early/2020/07/07/2020.02.24.961714.

Parker Hicks and Oluwatosin Oluwadare. HiCARN: resolution enhancement of Hi-C data using cascading residual networks. Bioinformatics, 38(9):2414–2421, 03 2022. ISSN 1367-4803. doi:10.1093/bioinformatics/btac156. URL https://doi.org/10.1093/bioinformatics/btac156.

Shanshan Zhang, Dylan Plummer, Leina Lu, Jian Cui, Wanying Xu, Miao Wang, Xiaoxiao Liu, Nachiketh Prabhakar, Jatin Shrinet, Divyaa Srinivasan, Peter Fraser, Yan Li, Jing Li, and Fulai Jin. Deeploop robustly maps chromatin interactions from sparse allele-resolved or single-cell hi-c data at kilobase resolution. Nature Genetics, 54(7): 1013–1025, July 2022a. ISSN 1546-1718. doi:10.1038/s41588-022-01116-w. URL https://www.nature.com/articles/s41588-022-01116-w.

Leina Lu, Xiaoxiao Liu, Wei-Kai Huang, Paola Giusti-Rodríguez, Jian Cui, Shanshan Zhang, Wanying Xu, Zhexing Wen, Shufeng Ma, Jonathan D. Rosen, Zheng Xu, Cynthia F. Bartels, Riki Kawaguchi, Ming Hu, Peter C. Scacheri, Zhili Rong, Yun Li, Patrick F. Sullivan, Hongjun Song, Guo-Li Ming, Yan Li, and Fulai Jin. Robust hi-c maps of enhancer-promoter interactions reveal the function of non-coding genome in neural development and diseases. Molecular Cell, 79(3):521–534.e15, August 2020. ISSN 1097-4164. doi:10.1016/j.molcel.2020.06.007.

Ruochi Zhang, Tianming Zhou, and Jian Ma. Multiscale and integrative single-cell hi-c analysis with higashi. Nature Biotechnology, 40(2):254–261, February 2022b. ISSN 1546-1696. doi:10.1038/s41587-021-01034-y. URL https://www.nature.com/articles/s41587-021-01034-y.

Ruochi Zhang, Tianming Zhou, and Jian Ma. Ultrafast and interpretable single-cell 3d genome analysis with fast-higashi. Cell Systems, 13(10):798–807.e6, October 2022c. ISSN 2405-4720. doi:10.1016/j.cels.2022.09.004.

Joshua D. Batson and Loic A. Royer. Noise2self: Blind denoising by self-supervision. ArXiv, abs/1901.11365, 2019.

Hanjun Shin, Yi Shi, Chao Dai, Harianto Tjong, Ke Gong, Frank Alber, and Xianghong Jasmine Zhou. Topdom: an efficient and deterministic method for identifying topological domains in genomes. Nucleic Acids Research, 44 (7):e70, Apr 2016. ISSN 0305-1048. doi:10.1093/nar/gkv1505. URL https://www.ncbi.nlm.nih.gov/pmc/articles/PMC4838359/.

Oana Ursu, Nathan Boley, Maryna Taranova, Y. X. Rachel Wang, Galip Gurkan Yardimci, William Stafford Noble, and Anshul Kundaje. Genomedisco: a concordance score for chromosome conformation capture experiments using random walks on contact map graphs. Bioinformatics (Oxford, England), 34(16):2701–2707, August 2018. ISSN 1367-4811. doi:10.1093/bioinformatics/bty164.

Suhas S.P. Rao, Miriam H. Huntley, Neva C. Durand, Elena K. Stamenova, Ivan D. Bochkov, James T. Robinson, Adrian L. Sanborn, Ido Machol, Arina D. Omer, Eric S. Lander, and Erez Lieberman Aiden. A 3d map of the human genome at kilobase resolution reveals principles of chromatin looping. Cell, 159(7):1665–1680, Dec 2014. ISSN 00928674. doi:10.1016/j.cell.2014.11.021. URL https://linkinghub.elsevier.com/retrieve/pii/S0092867414014974.

Merve Sahin, Wilfred Wong, Yingqian Zhan, Kinsey Van Deynze, Richard Koche, and Christina S. Leslie. Hic-dc+ enables systematic 3d interaction calls and differential analysis for hi-c and hichip. Nature Communications, 12 (1):3366, Jun 2021. ISSN 2041-1723. doi:10.1038/s41467-021-23749-x. URL https://www.nature.com/articles/s41467-021-23749-x.

Tsung-Han S. Hsieh, Claudia Cattoglio, Elena Slobodyanyuk, Anders S. Hansen, Oliver J. Rando, Robert Tjian, and Xavier Darzacq. Resolving the 3d landscape of transcription-linked mammalian chromatin folding. Molecular Cell, 78(3):539–553.e8, May 2020. ISSN 1097-4164. doi:10.1016/j.molcel.2020.03.002.

Matthew G. Heffel, Jingtian Zhou, Yi Zhang, Dong-Sung Lee, Kangcheng Hou, Oier Pastor-Alonso, Kevin D. Abuhanna, Joseph Galasso, Colin Kern, Chu-Yi Tai, Carlos Garcia-Padilla, Mahsa Nafisi, Yi Zhou, Anthony D. Schmitt, Terence Li, Maximilian Haeussler, Brittney Wick, Martin Jinye Zhang, Fangming Xie, Ryan S. Ziffra, Eran A. Mukamel, Eleazar Eskin, Tomasz J. Nowakowski, Jesse R. Dixon, Bogdan Pasaniuc, Joseph R. Ecker, Quan Zhu, Bogdan Bintu, Mercedes F. Paredes, and Chongyuan Luo. Temporally distinct 3d multi-omic dynamics in the developing human brain. Nature, page 1–9, October 2024. ISSN 1476-4687. doi:10.1038/s41586-024-08030-7. URL https://www.nature.com/articles/s41586-024-08030-7.

Zhou Wang, A.C. Bovik, H.R. Sheikh, and E.P. Simoncelli. Image quality assessment: from error visibility to structural similarity. IEEE Transactions on Image Processing, 13(4):600–612, 2004. doi:10.1109/TIP.2003.819861.

Neva C. Durand, Muhammad S. Shamim, Ido Machol, Suhas S.P. Rao, Miriam H. Huntley, Eric S. Lander, and Erez Lieberman Aiden. Juicer provides a one-click system for analyzing loop-resolution hi-c experiments. Cell Systems, 3(1):95–98, Jul 2016. ISSN 24054712. doi:10.1016/j.cels.2016.07.002. URL https://linkinghub.elsevier.com/retrieve/pii/S2405471216302198.

Sarah B. Reiff, Andrew J. Schroeder, Koray Kırlı, Andrea Cosolo, Clara Bakker, Luisa Mercado, Soohyun Lee, Alexander D. Veit, Alexander K. Balashov, Carl Vitzthum, William Ronchetti, Kent M. Pitman, Jeremy Johnson, Shannon R. Ehmsen, Peter Kerpedjiev, Nezar Abdennur, Maxim Imakaev, Serkan Utku Öztürk, Uğur Çamoğlu, Leonid A. Mirny, Nils Gehlenborg, Burak H. Alver, and Peter J. Park. The 4d nucleome data portal as a resource for searching and visualizing curated nucleomics data. Nature Communications, 13(1):2365, May 2022. ISSN 2041-1723. doi:10.1038/s41467-022-29697-4. URL https://www.nature.com/articles/s41467-022-29697-4.

Gökcen Eraslan, Lukas M. Simon, Maria Mircea, Nikola S. Mueller, and Fabian J. Theis. Single-cell RNA-seq denoising using a deep count autoencoder. Nature Communications, 10(1):390, January 2019. ISSN 2041-1723. doi:10.1038/s41467-018-07931-2. URL https://www.nature.com/articles/s41467-018-07931-2.

